# *Magnetospirillum magneticum* as a living iron chelator induces TfR1 upregulation and decreases cell viability in cancer cells

**DOI:** 10.1101/2020.05.28.121574

**Authors:** Stefano Menghini, Ping Shu Ho, Tinotenda Gwisai, Simone Schuerle

**Author notes:** These authors contributed equally (Stefano Menghini, Ping Shu Ho).

## Abstract

Interest has grown in harnessing biological agents for cancer treatment as dynamic vectors with enhanced tumor targeting. While bacterial traits such as proliferation in tumors, modulation of an immune response and local secretion of toxins have been well studied, less is known about bacteria as competitors for nutrients. Here, we investigated the use of a bacterial strain as a living iron chelator, competing for this nutrient vital to tumor growth and progression. We established an *in vitro* co-culture system consisting of the magnetotactic strain *Magnetospirillum magneticum* AMB-1 incubated under hypoxic conditions with human melanoma cells. Siderophore production by 10^8^ AMB-1/mL in human transferrin (Tf)-supplemented media was quantified and found to be equivalent to a concentration of 3.78 μM ± 0.117 μM deferoxamine, a potent drug used in iron chelation therapy. Our experiments revealed an increased expression of transferrin receptor 1 (TfR1) and a significant decrease of cancer cell viability, indicating the bacteria’s ability to alter iron homeostasis in human melanoma cells. Our results show the potential of a bacterial strain acting as a self-replicating iron-chelating agent, which could serve as an additional mechanism reinforcing current bacterial cancer therapies.

## 1. Introduction

Due to limited selectivity in systemically delivered cancer therapeutics, interest has grown in harnessing bacteria as living, tumor-targeting anticancer agents. The therapeutic potential of facultative anaerobic bacteria has been supported by studies demonstrating the delivery of non-pathogenic strains of *Escherichia coli* to solid flank tumors with associated tumor regression[1]. Additionally, safe administration of *Salmonella typhimurium* (VPN20009) has been shown for animal models and patients with metastatic melanoma [2,3]. Bacteria can act therapeutically by secreting innate or engineered toxins in situ (e.g. hemolysin E), transporting attached nanodrug formulations, or stimulating an immune response [4–7]. Colonizing bacteria can also engage in nutrient competition within the tumor microenvironment [8–10]. While the starvation of glucose as a crucial energy source to all cells has been studied intensively [11–13], other nutrients that are in specifically high demand by cancer cells might serve as more specific, vulnerable targets for deprivation.

Iron metabolism, for example, is significantly altered in mammalian tumor cells and recognized as a metabolic hallmark of cancer [14,15]. The main iron uptake mechanism adopted by most cells utilizes the internalization of transferrin receptor 1 (TfR1) upon binding of Fe (III)-bound transferrin (Tf). TfR1 expression positively correlates with cellular iron starvation and is upregulated in cancer cells, since malignant cells generally require a nutrient surplus [15–17]. Accordingly, several types of iron scavenging molecules have been utilized to compete with malignant cells for available iron sources and have demonstrated significant anti-neoplastic activity both, *in vitro* and *in vivo* [18–20] Promising bacteria-derived iron-chelating siderophores, such as deferoxamine (DFO), as well as synthetic iron chelators have been developed for therapeutic purposes [21]. However, non-negligible side effects, including systemic toxicity and low efficacy, have hampered their translation into clinical trials as therapeutic agents for cancer treatment [22–24].

Here we investigate the potential of a specific bacterial strain with high demand for iron to serve as local, self-replicating iron chelator that could thereby reduce systemic effects. Magnetotactic bacteria, like other bacteria, possess the ability to secrete high-affinity iron-scavenging siderophores. In particular, AMB-1 secrete both hydroxamate and catechol (3,4-dihydroxybenzoic acid) types of siderophores [25,26]. Unlike other bacteria, their demand for iron is particularly high, since this mineral is crucial both for their survival and synthesis of unique intracellular organelles called magnetosomes. [27,28]. These biomineralized magnetic nanocrystals are arranged in chains enclosed in a lipid bilayer and enable the bacteria to align with magnetic fields [29–31]. Furthermore, MTB are aerotactic, possessing an oxygen-sensing system that regulates motility in an oxygen gradient [32]. These features have previously been leveraged to magnetically guide MTB to the hypoxic core of solid tumors, yielding significantly higher tumor accumulation and penetration compared to their administration in the absence of external magnetic fields [33]. Once on site, nutrients from the tumor microenvironment are sourced to maintain proliferation and growth, and we hypothesize that MTB could induce iron deprivation of cancer cells.

To study this, we employed *Magnetospirillum magneticum* strain AMB-1 and first quantified the production of siderophores, benchmarked with molar concentrations of DFO. We then investigated the influence of AMB-1 on cell surface TfR1 expression using human melanoma cells and demonstrated the ability of AMB-1 to affect iron homeostasis. Finally, we examined the effect of AMB-1 on cancer cell growth *in vitro* by analyzing cell viability. The iron scavenging capabilities of bacterial strains with naturally high or enhanced siderophore production may act as an additional mechanism for bacterial cancer therapy, complementing or augmenting established bacterial anticancer mechanisms.

## 2. Results

### 2.1. AMB-1 produced siderophores affect human transferrin structure in mammalian cell culture medium

First, we sought to determine to what extent AMB-1 would produce siderophores in Dulbecco’s Modified Eagle’s medium (DMEM). Using the Chrome Azurol S (CAS) assay (Figure S1), 10^8^ AMB-1 bacteria were found to produce 0.10 ± 0.005 siderophore units in DMEM supplemented with 25 μM holo-transferrin (holo-Tf), while siderophore production in transferrin-free DMEM was negligible (Figure 1A). AMB-1 siderophore production was compared to the widely used iron chelator deferoxamine. It was found that the siderophores produced by 10^8^ AMB-1 in Tf-supplemented media was equivalent to 3.78 μM ± 0.117 μM deferoxamine (Figure 1B).

**Figure 1:**
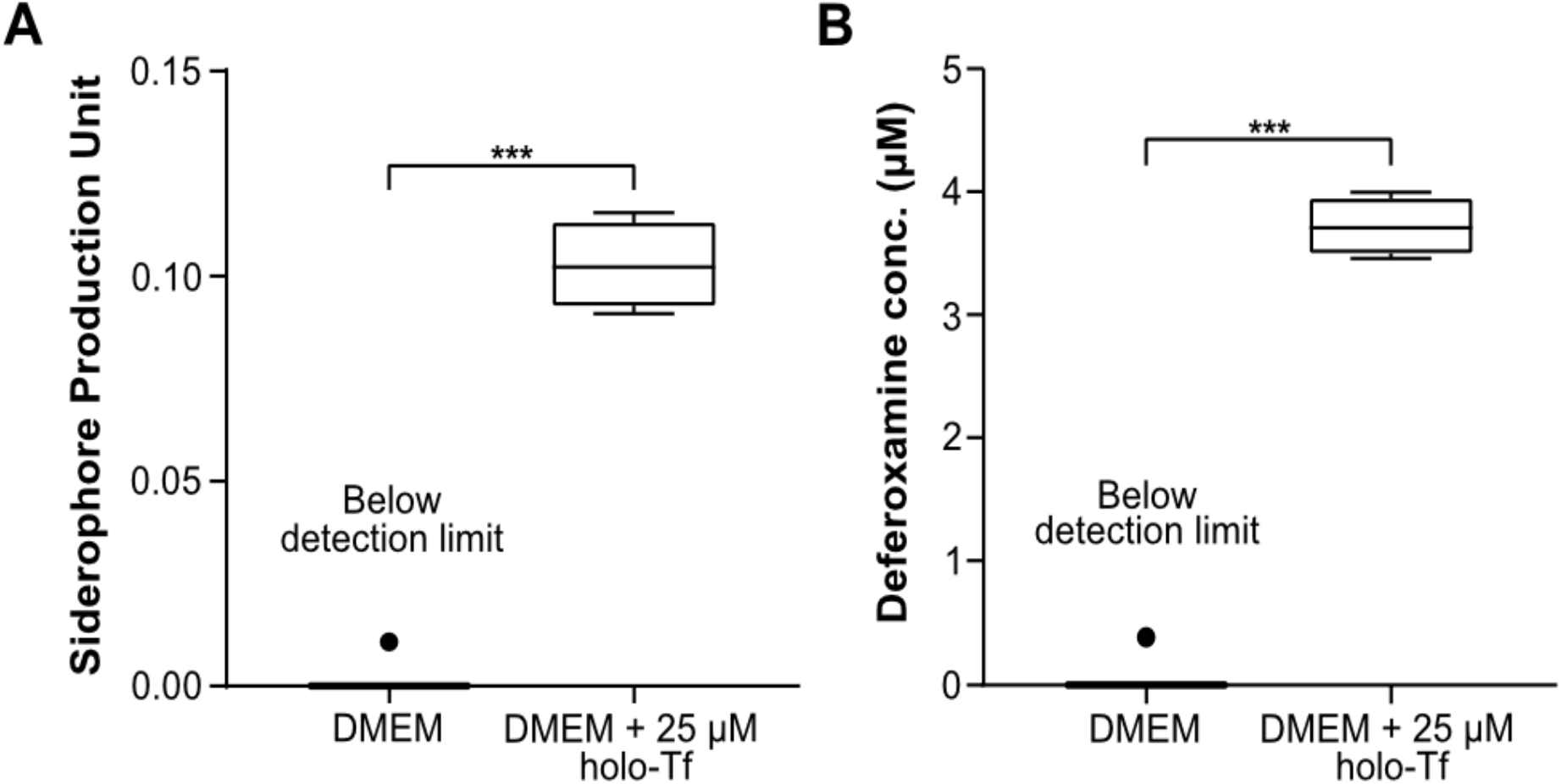
Quantification of siderophores produced by Magnetospirillum magneticum AMB-1 and analysis of their interaction with human transferrin. **(A)** Siderophores produced by AMB-1 were quantified by a chrome azurol S (CAS) assay in DMEM (condition 1) and DMEM supplemented with 25 μM holo-transferrin (condition 2) (n=4 per condition, statistical significance was assessed with an unpaired two-tailed t-test). **(B)** Siderophore production units plotted in terms of the inferred equivalent concentration of deferoxamine (n=4 per condition, statistical significance was assessed with an unpaired two-tailed t-test).

Having established the ability of AMB-1 to produce siderophores in Tf-supplemented media, we next determined whether AMB-1 would have an effect on human transferrin structure. SDS-PAGE analysis was used to compare DMEM supplemented with either iron-containing holo-Tf or iron-depleted apo-Tf. The apo-Tf appeared as a broader band on the SDS-gel compared to holo-Tf (Figure S2). Furthermore, we ascertained that holo-Tf structure was not affected by a 48 h incubation period at 30°C. To test whether the bacteria induced changes in Tf, AMB-1 were inoculated in DMEM supplemented with holo-Tf. This approach revealed that holo-Tf formed a broader band very similar to that seen for the apo-Tf band incubated in DMEM (Figure S2, lane 6). These experiments demonstrate that AMB-1 produced a quantifiable amount of siderophore when holo-Tf was supplemented to the mammalian cell culture media and that the structure of holo-Tf was affected by the bacteria.

### 2.2. AMB-1 upregulates TfR1 expression in human melanoma cells

To test whether AMB-1 can affect the iron uptake machinery in mammalian tumor cells we co-cultured AMB-1 with the human melanoma cell line MDA-MB-435S and monitored TfR1 expression using immunofluorescence. To mimic the tumor microenvironment, all experiments were performed under hypoxic conditions (Figure S3). The surface expression of TfR1 increased 2.7-fold on cancer cells co-cultured with live bacteria at AMB-1:MDA-MB-435S ratios as low as 10:1 (10^6^ AMB-1). The TfR1 upregulation was shown to increase with increasing bacteria ratios (Figure 2A, B). Deferoxamine was used here to create iron-deficient cell culture conditions as a positive control. MDA-MB-435S cells showed a significant and increasing upregulation of TfR1 surface expression up to 5.6-fold. To ensure that the upregulation of TfR1 expression was on the cell surface and not cytoplasmic, cell membrane integrity in the cultures was monitored. Less than 5% of cells were stained by the cell-impermeant DNA stain propidium iodide (PI), indicating cell membrane preservation over time (Figure 2C).

**Figure 2:**
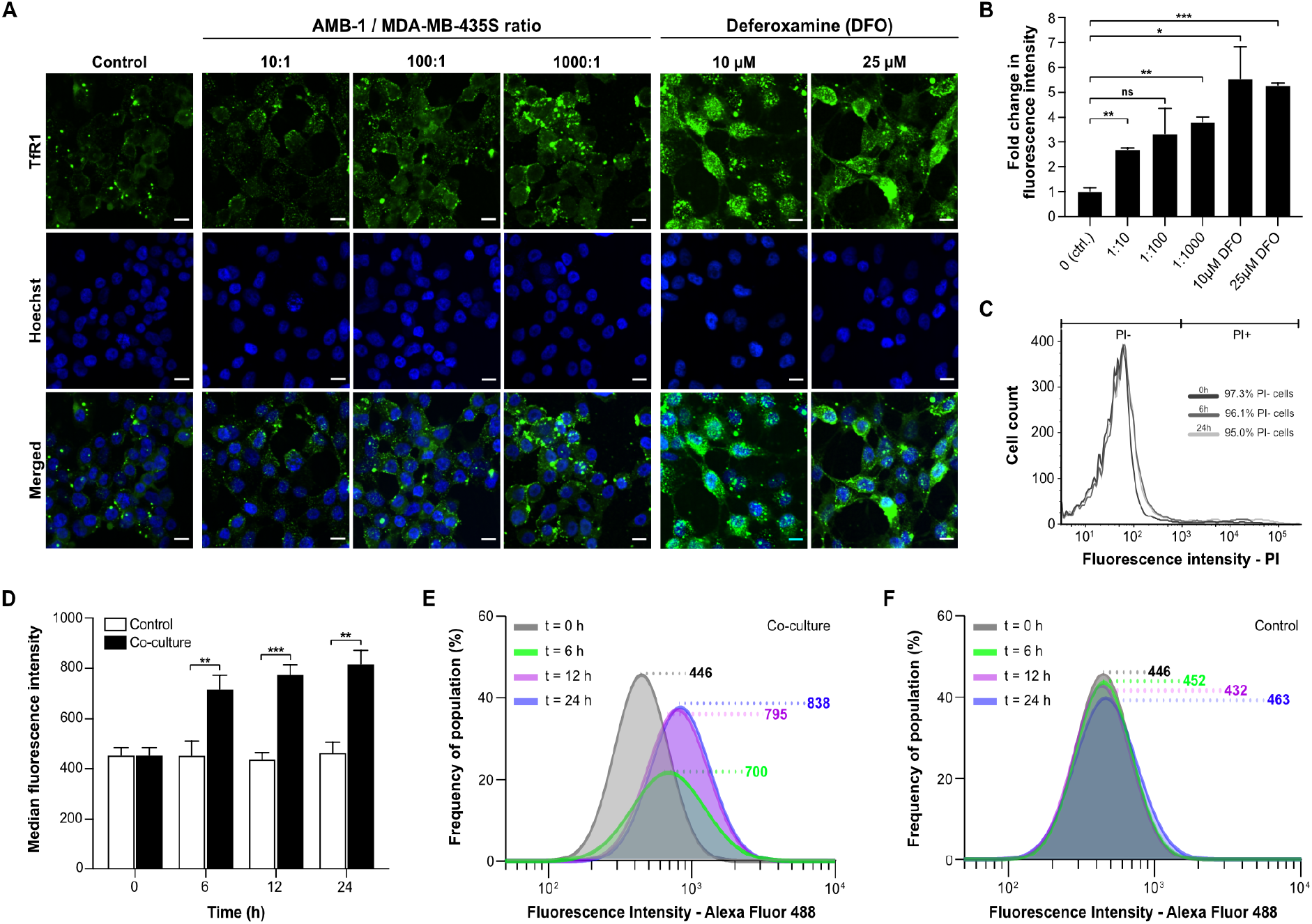
Analysis of TfR1 upregulation and cell surface expression on MDA-MB-435S. **(A)** Representative immunofluorescence images of human melanoma cells co-cultured under hypoxic conditions for 48 h with different ratios of AMB-1 bacteria and different concentrations of deferoxamine as a positive control. Images show MDA-MB-435S cells marked by anti-TfR1 antibody (green) and Hoechst 33342 (blue), (scale bar: 10 μM). **(B)** Quantification of the fold change in fluorescence intensity relative to the control condition, (n=2 biological replicates per condition, statistical significance was assessed with an unpaired two-tailed t-test). **(C)** Membrane integrity was measured as a graphical representation of PI negative and PI positive cell populations after 0, 6 and 24 h. **(D)** TfR1 median fluorescence intensity measured over 24 hours, (n=3 biological replicates per timepoint, statistical significance was assessed with an unpaired two-tailed t-test). **(E)** Representative log-normal fitted fluorescence intensity histograms of cell surface TfR1 expression on MDA-MB-435S cells in co-culture model and **(F)** negative control, (n=3 biological replicates per timepoint).

To gain insights on the TfR1 expression kinetics of the cell population, AMB-1-induced increase of cell surface TfR1 expression was analyzed over time. The effect, at an AMB-1:MDA-MB-435S ratio of 1000:1 was already apparent after 6 h of co-culture (Figure 1D). The fluorescent intensity after 24 h of co-culture was 1.8 times higher than the initial value, while the change reached 95% of the final value after 12 h (Figure 1E). Untreated cancer cells did not display any increase in fluorescence (Figure 1F). Upregulation of TfR1 could also not be found for non-magnetotactic bacteria with lower demand for iron, such as E. Coli [34–36]. When E. coli Nissle 1917 where incubated with melanoma cells at a ratio of of 1000:1, our highest bacteria to cell ratio tested for AMB-1, no detectable increase in TfR1 expression could be noted on the cell surface of MDA-MB-435S cells (Figure S4). Altogether, these findings show that the bactrial strain AMB-1 possesses a unique ability to induce TfR1 upregulation in the tested human melanoma cancer cell line, thereby suggesting a direct link between AMB-1 induced disruption of iron uptake and TfR1 expression.

### 2.3. Reduced viability of cancer cell lines upon incubation with AMB-1

Upon co-culturing melanoma cells with AMB-1 bacteria, we assessed cellular viability using MTT assay. A significant decrease in cell viability could be seen when cells were exposed to live bacteria at AMB-1:MDA-MB-435S ratios as low as 100:1 (10^7^ AMB-1). Incubation of MDA-MB-435S cells with bacteria (ratio 1000:1) resulted in an overall decrease of the mean cell viability of 62% (± 21.93%) (Figure 3). To ascertain that this effect was not restricted to one cell line, the experiment was repeated on an additional human cancer cell lines, MDA-MB-231. A significant decrease of 65% (± 23.37%) was detected in MDA-MB-231 cell viability once the cells where incubated with bacteria at a ratio of 1000:1. Supported by these observations, we showed that magnetotactic bacteria AMB-1 impact cancer cell viability, suggesting that they affect cancer cell growth *in vitro.*

**Figure 3:**
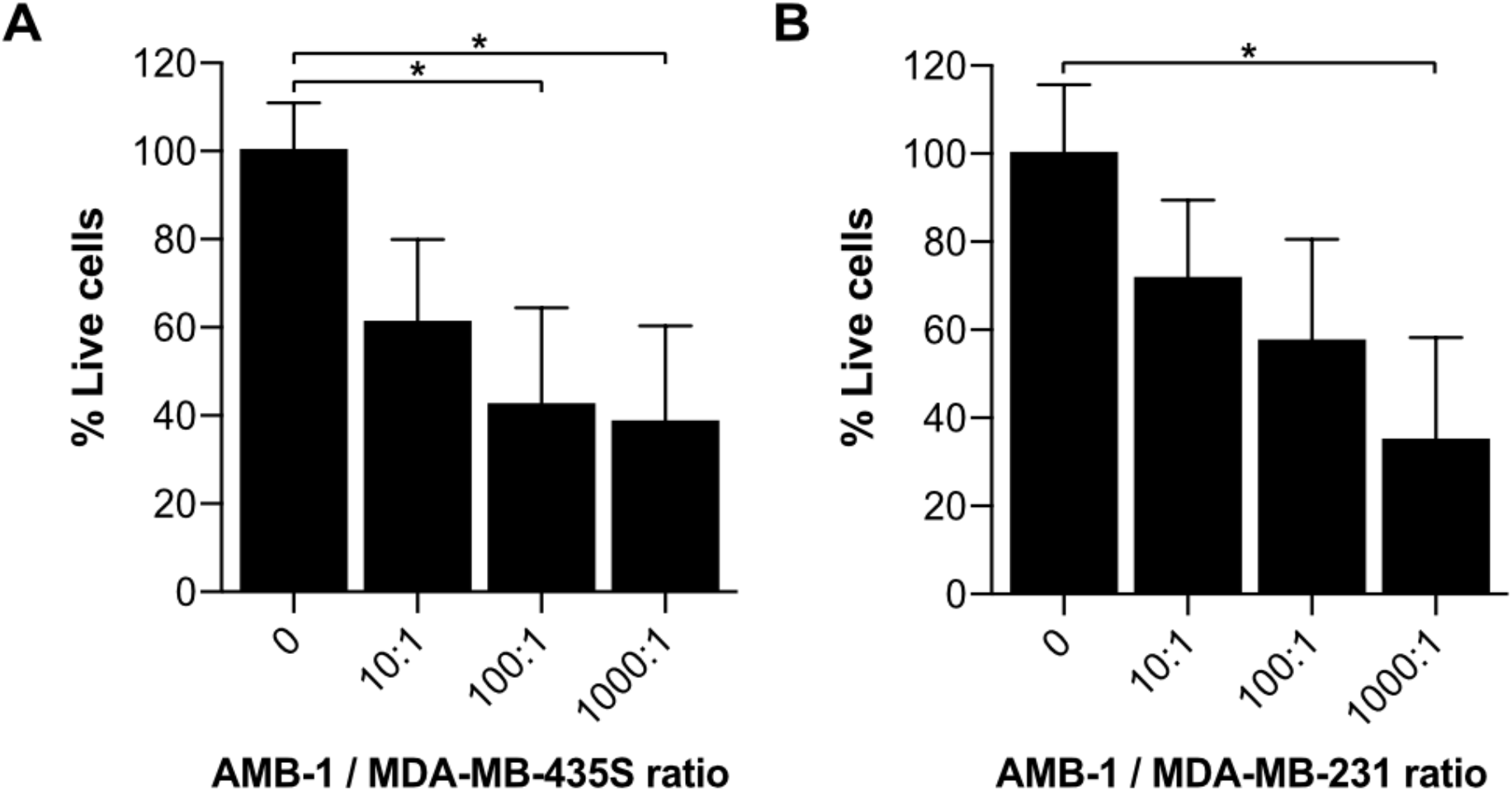
Investigation of cell growth upon incubation with AMB-1 bacteria. Cell viability of MDA-MB-435S and MDA-MB-231 was determined using an MTT assay and expressed as a percentage of the untreated cells. Viability (%) is expressed as mean ± SD of 3 individual biological replicates. Ordinary one-way ANOVA test was used to assess statistical significance.

## 3. Discussion

Magnetotactic bacteria acquire iron through siderophore-mediated uptake, as ferric and ferrous ions cannot directly enter bacteria cells. We quantified the number of siderophores produced by strain Magnetospirillum magneticum AMB-1 in mammalian cell culture medium and benchmarked the results with deferoxamine, a commonly used iron chelator. We also found that the bacteria affect human transferrin’s structure. The broadening of the holo-Tf band we observed in response to the addition of AMB-1 to the media suggests a change in molecular weight or conformation. Such a change in conformation would be consistent with the increased fraction of apo-Tf resulting from competition for binding of ferric iron with siderophores secreted by AMB-1. Comparing our findings to studies that showed the proteolytic cleavage of Tf by Prevotella nigrescens, we infer that specific degradation of the Tf by AMB-1 did not occur, since sub-products with lower molecular mass were not detected on the gel [37]. Therefore, our results suggest a loss of iron ions by holo-Tf, which is consistent with bacteria-produced siderophores having a higher affinity for Fe ions compared to human transferrin [38,39]. This higher affinity could be exploited by AMB-1 to efficiently compete for ferric ions with the host cells, resulting in iron starvation for the latter.

We then showed that AMB-1 inoculation with human melanoma cell cultures affects iron homeostasis of the cancer cells. Iron homeostasis is essential for normal cell growth and development, and iron starvation is mainly characterized by alterations in the iron import machinery, specifically by an upregulation of the transferrin receptor 1 on the cell surface. Increased TfR1 expression found on MDA-MB-435S melanoma cancer cells correlates with increasing bacteria ratios. This finding suggests that AMB-1 effectively competes for free iron ions and therefore limits the mineral’s availability to MDA-MB-435S cells (Figure 2A, B). Moreover, a significant increase of TfR1 expression could already be detected 6 h after inoculation (Figure 2D-F). Similarly, the cancer cells showed a significant upregulation of TfR1 surface expression after incubation with deferoxamine (10 μM and 25 μM), in line with previous reports on cellular iron deficiency [15,16,40]. These observations demonstrated that AMB-1 affects the iron import mechanisms of human melanoma cells, acting as an effective competitor for iron when in co-culture with MDA-MB-435S cells.

Furthermore, we assessed the impact of AMB-1 cells on cancer cell growth (Figure 3). Earlier studies indicated the benefits of adding iron chelators to cancer cells, showing a reduction of cell growth upon treatment [19,24,40]. Our results confirmed that an increasing number of AMB-1 added to co-culture correlated with a decrease in the percentage of viable cancer cells. This effect could be detected on cancer cells from two different lineages. At the highest investigated bacteria to cell ratio (1000:1), the melanoma cells MDA-MB-435S displayed an overall viability of 38% and the viability of breast cancer cells MDA-MB-231 was 32%. These findings suggest that the high requirement for iron exhibited by MTB causes them to actively compete for Fe (III) with cancer cells, leading to a nutrient shortage associated with reduced viability.

Our data support the idea that AMB-1 have the ability to act as living iron chelators by secreting a quantifiable amount of siderophores. We showed that 10^8^ AMB-1/mL can produce high-affinity iron scavenging molecules equivalent to 3.78 μM deferoxamine over 24 h (Figure 1B). Previous works demonstrated that the treatment of different cell lines with 10 μM - 30 μM deferoxamine significantly reduced cell viability *in vitro* [19,40]. Moreover, a significant diminution of cell viability was even detected at the lower deferoxamine concentration of 2.5 μM when combined with the chemotherapeutic drug cisplatin [19]. Nonetheless, the implementation of molecular iron scavenging molecules in translational medicine is hampered by elevated systemic toxicity, as well as limited tumor selectivity. These challenges might be overcome by implementing bacteria as direct competitors for nutrients at the tumor site. Several approaches have been investigated in the past to deliver magnetotactic bacteria to solid tumors. For example, intravenously introduced AMB-1 have been shown to colonize tumor xenografts 3-6 days after injection [41]. More recent attempts describe the use of peritumoral injection, followed by magnetic guidance [33] as well as the powered actuation of swarms of AMB-1 with a rotating magnetic field [42]. Information on immune response in these studies is limited, though intratumoral injection of chains of magnetosomes coated with endotoxins has been shown to elicit recruitment of immune cells to the tumor site and trigger cancer regression [43]. Previous studies have also reported of the ability of magnetotactic bacteria to reproduce and form magnetite when cultured at 37 degrees [41], a finding we independently corroborated by investigating AMB-1 proliferation and viability over 48 hours at 37 °C (Figure S5). Furthermore, the intrinsic magneto-aerotactic capability of MTB allows them to regulate their motility towards environments with low oxygen concentration as well as react to externally applied magnetic fields [31,32]. Aerotaxis and anaerobic traits have also been leveraged in other strains, such as *Salmonella*, enabling them to act as bacterial anti-cancer agents that target necrotic tumor microenvironments with poor oxygen supply [44–46]. Overall, the intrinsic abilities of AMB-1 to self-replicate, respond to magnetic fields, and secrete sustained doses of siderophores warrants further study in the context of cancer therapy. By combining the benefits of bacterial cancer therapy with the iron chelation and other traits of AMB-1, we envision that magnetotactic bacteria could become a valid therapeutic agent to implement against cancer.

Our work motivates the use of living AMB-1 as self-replicating iron scavenging organisms actively competing for this vital nutrient, with the possibility of compromising the survival of cancer cells. Further application could include the use of tumor-targeting organisms both as a monotherapy and as a combination therapy with established anti-neoplastic drugs to obtain optimal clinical outcomes. Moreover, the unique characteristics of magnetotactic bacteria could be exploited to engineer iron-scavenging strains of surrogate commensal and attenuated bacteria that have already been established as anti-cancer agents [3,7]. This work lays the foundation for future investigations which combine iron chelation with bacterial cancer therapy to enhance existing therapeutic strategies and open new frontiers for combating cancer.

## 4. Materials and Methods

### 4.1. Bacterial strain and culture condition

*Magnetospirillum magneticum* AMB-1, a strain of magnetotactic bacteria, was purchased from ATCC (ATCC, Manassas, Virginia, USA). AMB-1 bacteria were grown anaerobically at 30°C, passaged weekly and cultured in liquid growth medium (ATCC medium: 1653 Revised Magnetic Spirillum Growth Medium). *Magnetospirillum magneticum* Growth Media (MSGM) contained the following per liter: 5.0 mL Wolfe’s mineral solution (ATCC, Manassas, Virginia, USA), 0.45 mL Resazurin, 0.68 g of monopotassium phosphate, 0.12 g of sodium nitrate, 0.035 g of ascorbic acid, 0.37 g of tartaric acid, 0.37 g of succinic acid and 0.05 sodium acetate. The pH of the media was adjusted to 6.75 with sodium hydroxide (NaOH) and then sterilized by autoclaving at 121°C. 10 mM ferric quinate (200x) Wolfe’s Vitamin Solution (100x) (ATCC, Manassas, Virginia, USA) were added to the culture media shortly before use. The concentration of AMB-1 in solution was determined by optical density measurement (Spark, Tecan, Männedorf, Switzerland) and the approximate number of bacteria was extrapolated from a standard curve.

Start of *E. coli* cultures were achieved by picking single colonies from LB agar plates and subsequent inoculation in liquid LB media. Pre-culture of bacteria were performed the day before the experiment in liquid LB media over night at 37°C on a shaking device. On the day of the experiment, an approximate number of E. coli in solution was then determined by optical density measurement (Spark, Tecan, Männedorf, Switzerland),

### 4.2. CAS assay to asses siderophore quantification

*Magnetospirillum magneticum* AMB-1 were cultured in 1.7 mL phenol red-free DMEM (11054020, Invitrogen, Carlsbad, California, USA) supplemented with GlutaMAX (35050061, Invitrogen, Carlsbad, California, USA) in a sealed 1.5 mL Eppendorf tube at 37°C for 24 h. Fetal bovine serum (FBS, Biowest, Nuaille, France) was excluded from the media and replaced with a known concentration of iron source; 25 μM holo-transferrin (T0665, Sigma-Aldrich, St. Louis, Missouri, USA). Quantification of siderophores produced by AMB-1 was performed using the Chrome Azurol S (CAS) assay (199532, Sigma-Aldrich, St. Louis, Missouri, USA) [47]. 100 μL of each sample’s supernatant was collected and mixed with 100 μL CAS assay solution on a transparent 96-well plate. The assay was then incubated in the dark at room temperature for 1 h before the absorbance was measured at 630 nm on a multimode microplate reader (Spark, Tecan, Männedorf, Switzerland). The measurement was expressed in siderophore production unit (s.p.u.), which was calculated as follows:

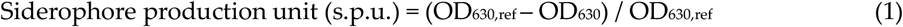

DMEM supplemented with different concentrations of deferoxamine mesylate salt (DFO, D9533, Sigma-Aldrich, St. Louis, Missouri, USA) was prepared by serial dilution and used to generate a calibration curve (Figure S1).

### 4.3. Analysis of human transferrin using SDS-PAGE Electrophoresis

AMB-1 bacteria (1 × 10^8^ cells/mL) were cultured in 1.7 mL phenol red-free DMEM (11054020, Invitrogen, Carlsbad, California, USA) in a sealed 1.5 mL Eppendorf tube at 30°C for 48 h. Excess volume was used to ensure no or minimal air was trapped in the tubes. 25 μM holo-transferrin (holo-Tf, T4132, Sigma-Aldrich, St. Louis, Missouri, USA), or 25 μM apo-transferrin (apo-Tf, T2036, Sigma-Aldrich, St. Louis, Missouri, USA) respectively were added to the mammalian cell culture media. Changes in transferrin molecular mass during the growth of AMB-1 were evaluated by sodium dodecyl sulfate-polyacrylamide gel electrophoresis (SDS-PAGE) analysis of culture supernatant. Electrophoresis was conducted using the protocol described by Laemmli [48] and protein loading of each sample was normalized to 2 μg. Proteins were visualized using SYPRO ruby protein stain (1703126, Bio-rad, Hercules, California, USA). The electrophoresis chamber and the reagents were purchased from Bio-rad. Stained gels were imaged using a fluorescent scanner (Sapphire Biomolecular Imager, Azure Biosystems, Dublin, California, USA) at 488 nm excitation and 658 nm emission.

### 4.4. Mammalian cell culture

Human melanoma MDA-MB-435S cells (ATCC, Manassas, Virginia, USA) were cultured in high glucose Dulbecco’s Modified Eagle’s Medium (DMEM, Invitrogen, Carlsbad, California, USA) supplemented with 10% fetal bovine serum (FBS, Biowest, Nuaille, France) and 1% penicillin-streptomycin (CellGro, Corning, New York, USA). All cells were incubated at 37°C in a humidified atmosphere with 5% CO_2_ at 37°C.

### 4.5. Co-culture of mammalian cancer cells with magnetotactic bacteria

Human melanoma MDA-MB-435 cells (1 × 10^5^ cells) were cultured on 12-well plates and incubated in a 5% CO_2_ incubator at 37°C for 24 h. For microscopic analysis at high magnification (> 40x), a circular cover slip was placed in each well prior to cell seeding. Following incubation, *Magnetospirillum magneticum* AMB-1 (1 × 10^6^ – 1 × 10^8^ cells) were introduced into the wells. The well plate was stored in a sealable bag and the bag was flushed with nitrogen for 15 min in order to produce hypoxic conditions. The setup with the 12-well plate was then incubated at 37°C for 48 h. To serve as negative and positive controls, 0, 10 μM and 25 μM of the iron-chelating agent deferoxamine mesylate (D9533, Sigma-Aldrich, St. Louis, Missouri, USA) was added to the MDA-MB-435S cell culture in place of AMB-1 bacteria.

### 4.6. Immunofluorescence labelling of MDA-MB-435S cells

After the co-culture, cells were washed with ice cold 1X Dulbecco’s Phosphate-Buffered Saline solution (DPBS, Gibco, Carlsbad, California, USA) and then blocked with a 1% Bovine Serum Albumin (BSA, Sigma-Aldrich, St. Louis, Missouri, USA) solution diluted in 1X DPBS. The cells were then incubated with 10 μg/mL primary anti-TfR1 antibody (ab84036, Abcam, Cambridge, UK) on ice in dark for one h. Subsequently, the cells were washed with ice-cold DPBS and incubated with 20 μg/mL secondary goat anti-rabbit antibody (ab150077, Abcam, Cambridge, UK) and 25 μg/mL Hoechst 33342 (H3570, Thermo Fisher Scientific, Waltham, Massachusetts, USA) on ice in dark for another hour. Next, the cells were washed with ice-cold 1X PBS twice and fixed with a 2% paraformaldehyde (PFA) solution. Fixed cells were washed three times with 1X DPBS and the cover slips were mounted on glass slides and stored overnight in dark at 4°C. A Nikon Eclipse Ti2 microscope equipped with a Yokogawa CSU-W1 Confocal Scanner Unit and Hamamatsu C13440-20CU ORCA Flash 4.0 V3 Digital CMOS camera were used for visualization. Microscope operation and image acquisition was performed using Nikon NIS-Elements Advanced Research 5.02 (Build 1266) software. ImageJ v2.0 (NIH) was used to process the obtained images.

### 4.7. Evaluation of fluorescently labelled MDA-MB-435S cells by flow cytometry

Flow cytometry was used to measure the expression of fluorescently labelled TfR1 on the surface of MDA-MB-435S cells. Cells were harvested at different timepoints during co-culture (0h, 6h, 12h, 24h) and washed in cold 1X DPBS (Gibco Carlsbad, California, USA). Harvested cells were stained with primary anti-TfR1 antibody (ab84036, Abcam, Cambridge, UK) at a concentration of 10 μg/mL. After 1 h of incubation on ice, cells were washed twice with 1X DPBS and then stained with 20 μg/mL secondary goat anti-rabbit antibody (ab150077, Abcam, Cambridge, UK). Finally, cells were washed twice with 1X DPBS and analyzed by flow cytometry with BD LSRFortessa (BD Biosciences, San Jose, California, USA) using a 488nm excitation laser and 530/30 and 690/50 band pass emission filters for detection. FlowJo™ (Tree Star) software was used to evaluate the data.

Flow cytometry was used to assess the cell membrane integrity of MDA-MB-435S cells. Cells were harvested at different timepoints during co-culture (0h, 6h, 24h) and washed in cold 1X DPBS. Collected cells were stained with 1 μg/mL Propidium Iodide (V13242, Thermo Fisher Scientific, Waltham, Massachusetts, USA) and incubated for 30 in a humidified atmosphere with 5% CO_2_ at 37°C. Finally, cells were washed twice with 1X DPBS and analyzed by flow cytometry with BD LSRFortessa (BD Biosciences, San Jose, California, USA) using a 488nm excitation laser and 610/10 bandpass emission filters for detection. FlowJo™ (Tree Star) software was used to evaluate data and graphs were plotted using Prism 8.0 (GraphPad).

### 4.8. Investigation of cell viability using MTT assay

CyQUANT MTT Cell Viability Assay (V13154, Thermo Fisher Scientific, Waltham, Massachusetts, USA) was used to measure the viability of human melanoma cells MDA-MB-435S and human breast cancer cell lines MDA-MB-231. Cells were plated in 96 well culture plates (50000 cells/well) and incubated in a 5% CO_2_ incubator at 37°C for 24 h. Co-culture was then performed by adding AMB-1 bacteria at different ratios (10:1, 100:1, 1000:1) under hypoxic conditions, as described earlier. After 24 h of incubation, cells were washed 3x with cold DPBS (Gibco Carlsbad, California, USA) to remove the bacteria. Next, 100μL of DMEM and 10μL of MTT stock solution (12 mM) were added to the wells and cells were then incubated at 37°C for 4 hours. The media was removed, and Formazan crystals formed by the cells were dissolved in 50 μL DMSO. The absorbance was measured at 540 nm using a multimode microplate reader (Spark, Tecan, Männedorf, Switzerland). Background signal was subtracted from the final values and data was first normalized to an untreated control and then plotted as a percentage of the untreated cells.

### 4.9. Staining and quantification of AMB-1 bacteria

AMB-1 bacteria were incubated in phenol red-free DMEM (21063-029, Gibco Carlsbad, California, USA) at 37°C and analyzed at different intervals (Day 0, Day 1, and Day 2). At each timepoint bacteria were collected, stained using BacLight viability Kit (L7007, BacLight Bacterial Viability Kit, Thermo Fisher Scientific, Waltham, Massachusetts, USA), and incubated in the dark for 15 min. Visualization and image acquisition was performed using confocal microscopy (Nikon Eclipse Ti2). ImageJ v2.0 (NIH) was used to process the obtained images. Quantification of bacteria was performed using a particle counting system, Multisizer 4e Coulter Counter (Beckman Coulter, Brea, California, USA).

### 4.10. Statistics and data analysis

All graphs and statistical analyses were generated using Prism 8.0 (GraphPad). Statistical significance and number of replicates of the experiments are described in each figure and figure legend. Error bars, where present, indicate the standard error of the mean (SD). P values are categorized as * P<0.05, ** P<0.01, and *** P<0.001.

## Author contributions

SS and PSH conceived and designed the experiments. PSH and SM collected and analyzed data. SM wrote the manuscript. TG assisted the authors and contributed to the paper revisions. SS supervised the study and helped writing the paper.

## Acknowledgements

The authors thank Cameron Moshfegh for helpful discussions. Technical support was provided by the Flow Cytometry Core Facility at ETH Zurich for flow cytometry measurements. The authors appreciate the help of Nima Mirkhani with AMB-1 cultures. The authors thank Guy Riddihough (Life Science Editors) for his editing support and Michael G. Christiansen for critically reviewing the manuscript and for his assistance in creating the hypoxia device. The authors would finally like to extend their gratitude to Dr. Tim Keys and Prof. Emma Slack’s group for kindly providing the *E. coli* Nissle 1917. A preprint of this manuscript is available at bioRxiv, MS ID#:BIORXIV/2020/121574.

## Abbreviations

Tf: Transferrin
TfR1: Transferrin receptor 1
MTB: Magnetotactic bacteria
DFO: Deferoxamine
DMEM: Dulbecco’s Modified Eagle’s medium

