## Supplementary Material for "*Magnetospirillum magneticum* as a living iron chelator induces TfR1 upregulation and decreases cell viability in cancer cells"

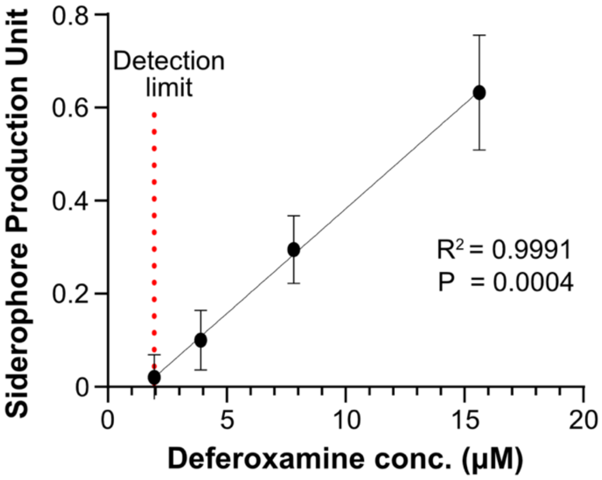


**Figure S1:** Calibration curve representing the siderophore production unit plotted against the concentration of deferoxamine (n= 3).

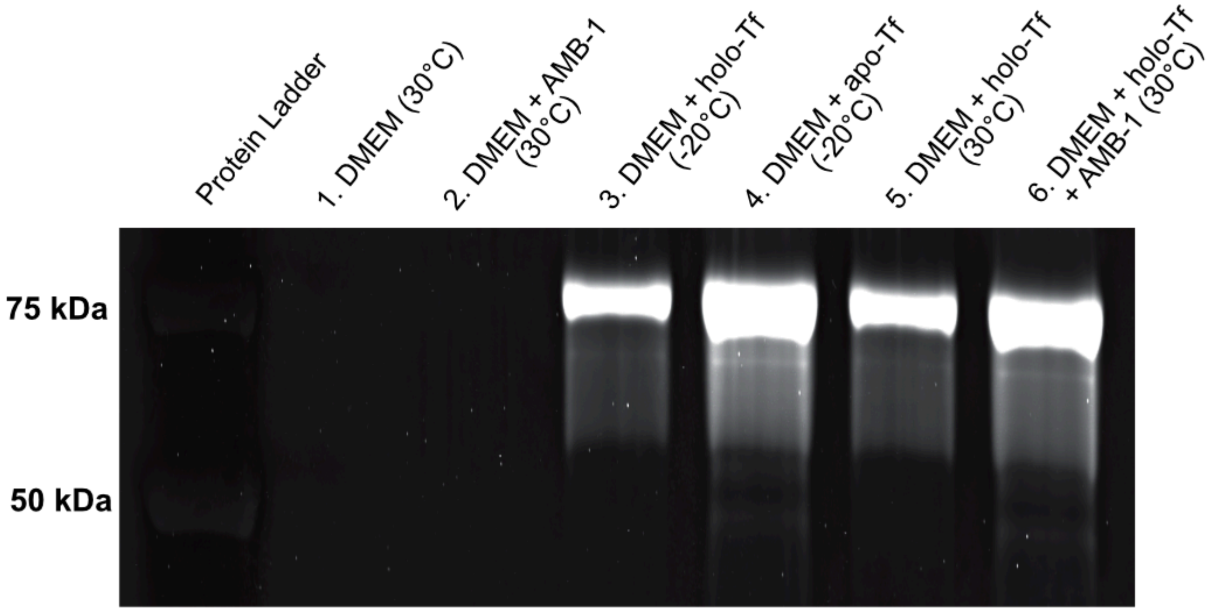


**Figure S2:** SDS-PAGE analysis displaying the effect of AMB-1 on the structure of human transferrin. Tested conditions are indicated in the figure, with holo-Tf corresponding to saturated transferrin and apo-Tf corresponding to non-saturated transferrin.

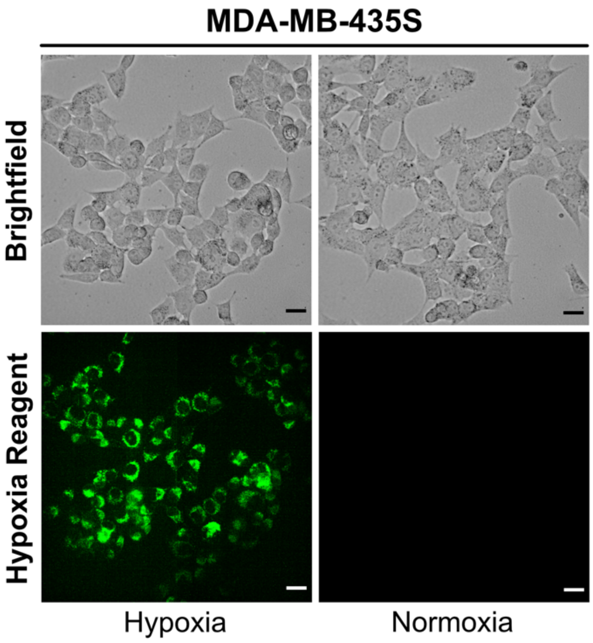


**Figure S3:** Comparison of in vitro cancer cell culture under either hypoxic or normoxic conditions. Representative fluorescence and brightfield images of MDA-MB-435S cells stained with Image-IT Green Hypoxia Reagent (green), (scale bar: 25 μM).

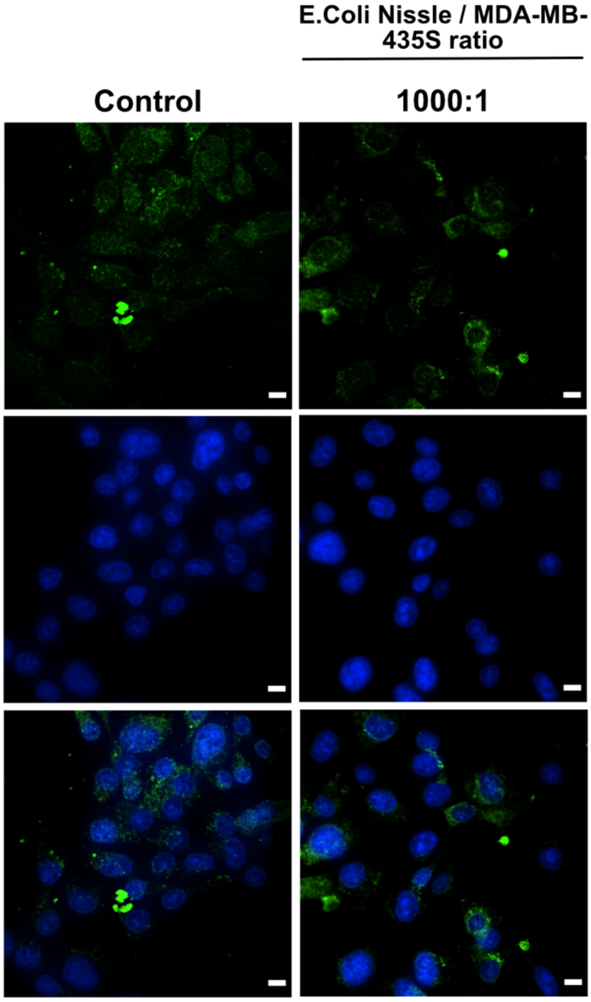


**Figure S4:** Immunofluorescence images of human melanoma cells co-cultured under hypoxic conditions for 48 hours with E. coli Nissle 1917. Images show MDA-MB-435S cells marked by anti-TfR1 antibody (green) and Hoechst 33342 (blue), (scale bar: 10 μM).

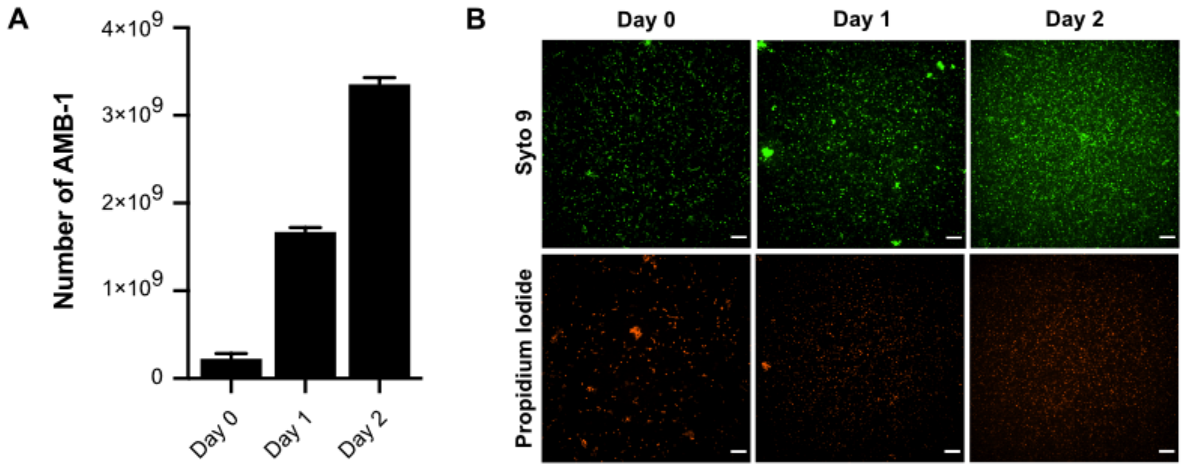


**Figure S5:** Quantification and visualization of proliferating AMB-1 bacteria. **(A)** Graphical representation of the increasing number of AMB-1 bacteria measured at different timepoints over the course of 2 days, (n=3, technical replicates). Counting of bacteria was performed using a Multisizer 4e (Coulter Counter). **(B)** Representative fluorescence images show AMB-1 bacteria stained with Syto 9 (green) and Propidium Iodide (red). Bacteria were imaged at different timepoints (Day 0, Day 1, and Day 2), demonstrating the increase of viable cells (scale bar: 25 µM).
